# Clathrin Senses Membrane Curvature

**DOI:** 10.1101/2020.06.04.134080

**Authors:** Wade F. Zeno, Jacob B. Hochfelder, Ajay S. Thatte, Liping Wang, Avinash K. Gadok, Carl C. Hayden, Eileen M. Lafer, Jeanne C. Stachowiak

## Abstract

The ability of proteins to sense membrane curvature is essential to diverse membrane remodeling processes including clathrin-mediated endocytosis. Multiple adaptor proteins within the clathrin pathway have been shown to assemble together at curved membrane sites, leading to local recruitment of the clathrin coat. Because clathrin does not bind to the membrane directly, it has remained unclear whether clathrin plays an active role in sensing curvature or is passively recruited by its adaptor proteins. Using a synthetic tag to assemble clathrin directly on membrane surfaces, here we show that clathrin is a strong sensor of membrane curvature, comparable to previously studied adaptor proteins. Interestingly, this sensitivity arises from clathrin assembly, rather than from the properties of unassembled triskelia, suggesting that triskelia have preferred angles of interaction, as predicted by earlier structural data. Further, when clathrin is recruited by adaptors, its curvature sensitivity is amplified by two to ten-fold, such that the resulting protein complex is up to 100 times more likely to assemble on a highly curved surface, compared to a flatter one. This exquisite sensitivity points to a synergistic relationship between the coat and its adaptor proteins, which enables clathrin to pinpoint sites of high membrane curvature, an essential step in ensuring robust membrane traffic. More broadly, these findings suggest that protein networks, rather than individual protein domains, are likely the critical drivers of membrane curvature sensing.

## INTRODUCTION

Clathrin-mediated endocytosis is the best understood cellular mechanism for internalization of membrane proteins, lipids, and extracellular cargo^1, 2^. Because the clathrin coat does not bind directly to lipids and cargo proteins, clathrin triskelia must be recruited to the membrane surface by adaptor proteins^3^. During assembly of a clathrin-coated vesicle, dozens of adaptors come together at sites of high membrane curvature, where they recruit triskelia, driving assembly of the clathrin coat^4, 5^. Many clathrin adaptor proteins have been shown to bind preferentially to highly curved membrane surfaces^6^. Some of these adaptors, such as Amphiphysin, Endophilin, and Fcho, contain inherently curved BAR (bin/amphiphysin/Rvs) family domains^7^. These domains are thought to sense membrane curvature through a scaffolding mechanism in which the curved membrane binding surface is more strongly attracted to membranes that match its curvature^8^. In contrast other adaptors such as Epsin^9^ and AP180/Calm^10^, include amphipathic helices or lipophilic motifs that can insert between membrane lipids. These wedge-like motifs are thought to sense the presence of gaps between lipid head groups, which are more abundant on highly curved membrane surfaces^11^. Additionally, intrinsically disordered regions, which are found in most adaptor proteins^3, 12^, are potent sensors of membrane curvature through a combination of entropic^13^ and electrostatic mechanisms^14^.

While the curvature sensing properties of adaptor proteins are increasingly well understood, it has remained unclear to what extent the clathrin coat itself can assemble preferentially at curved membrane sites. Based on electron micrographs of clathrin coated buds in cells^15^, as well as the high resolution structures of minimal clathrin baskets^16^, it is clear that clathrin triskelia are capable of assembling to form highly curved surfaces. These observations suggest, if indirectly, that clathrin assembly favors high curvature. However, flat clathrin lattices have frequently been observed in cells^15^, and recently under physiological conditions^17^. Therefore, it is presently unclear whether clathrin prefers to assemble into curved or flat lattices.

The past few years have seen a resurgence of the debate surrounding the relationship between clathrin assembly and membrane curvature. While earlier work indicated that the curvature of a clathrin coated vesicle remains approximately constant during its development^18^, recent reports have suggested that clathrin initially assembles into a relatively flat lattice and then makes an abrupt transition to a highly curved morphology^19,20^. To better understand these conflicting observations, it is important to determine the impact of membrane curvature on the assembly of the clathrin lattice. Toward this goal, here we directly measure the partitioning of clathrin among membrane surfaces that span a broad range of curvatures. To isolate the curvature sensitivity of clathrin from that of its adaptor proteins, we use a recombinant form of clathrin that contains an N-terminal histidine tag. This tag can recruit clathrin to membrane surfaces that contain synthetic, histidine-binding lipids. Importantly, the location of the tag, near the clathrin N-terminal domain, mimics the natural orientation of clathrin with respect to the membrane surface^16^. In this way, assembly of the clathrin lattice occurs directly on the membrane, without the requirement for an adaptor protein. Using this approach, our data reveal that clathrin strongly prefers to assemble on curved membrane surfaces, displaying a sensitivity to membrane curvature that is on par with that of many clathrin adaptor proteins. Importantly, this preference is lost when clathrin assembly is inhibited by high pH, suggesting that curvature sensing by clathrin arises from a preferred orientation of assembly, rather than from the properties of individual triskelia. Interestingly, when curvature sensitive adaptor proteins, such as epsin1 and amphiphysin1, are used to recruit clathrin, the overall sensitivity of the membrane-bound protein complex increases dramatically. These results suggest that adaptors and clathrin play synergistic roles in driving robust coated vesicle assembly at curved membrane sites.

## RESULTS

### Curvature sensitivity of clathrin in the absence of adaptor proteins

Clathrin assembles into lattices that consist of pentagonal and hexagonal facets^21^, Figure 1**a**. In the cell, these lattices are formed on the surfaces of intracellular membranes, when clathrin is recruited by adaptor proteins. However, *in vitro* experiments have revealed that concentrated solutions of clathrin alone can assemble into spherical, cage-like lattices, illustrating that clathrin assembly does not strictly require adaptors^22, 23^, To examine clathrin-membrane interactions directly, we expressed and purified recombinant clathrin with a hexa-histidine tag on the terminal domain of the heavy chain (his-clathrin). We have previously demonstrated that this recombinant clathrin assembles into uniform cages that are identical to those formed by clathrin isolated from bovine brain^24^. By using lipids with functionalized Ni^2+^ headgroups, we were able to examine clathrin-membrane interactions in the absence of adaptor proteins.

**Figure 1.**
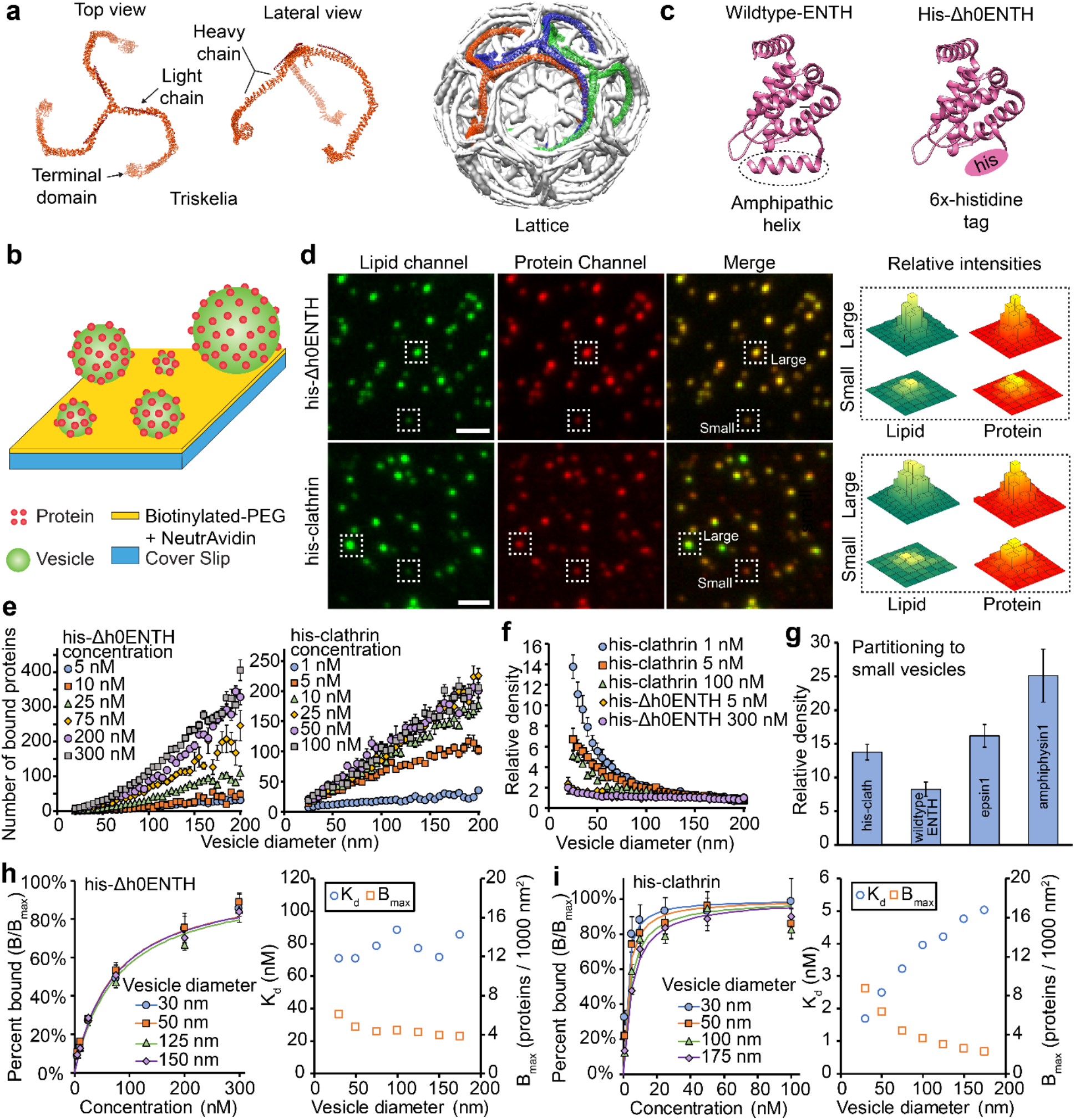
Histidine-tagged clathrin binds preferentially to vesicles with small diameters. **(a)** Structures of clathrin as a single triskelion and assembled into a basket (Protein Data Bank 3IYV). **(b)** Schematic of the assay used to tether vesicles and monitor protein binding. **(c)** Structures of wildtype-ENTH and his-Δh0ENTH (Protein Data Bank 1H0A). **(d)** Representative fluorescence images of tethered vesicles and the proteins that were bound to them. Vesicles were fluorescently labeled with ATTO 465-DOPE. Proteins were fluorescently labeled with Alexa Fluor 647. Protein concentrations used for these images were 100 nM for his-Δh0ENTH and 25 nM for his-clathrin. Scale bars represent a distance of 2 μm. **(e)** Number of proteins bound to vesicles that were exposed to either his-Δh0ENTH or his-clathrin. **(f)** Relative densities of his-Δh0ENTH and his-clathrin among vesicles with different diameters. These densities are normalized by the average value for vesicles between 160 and 200 nm. **(g)** Comparision of his-clathrin’s curvature sensitivity in **f** to previously measured values^13^ for ENTH, epsin1, and amphiphysin1. **(h)** Binding isotherm for his-Δh0ENTH with the corresponding lines of best fit from equation 1 and the resulting best-fit parameters. **(i)** Binding isotherm for his-clathrin with the corresponding lines of best fit from equation 1 and the resulting best-fit parameters. All vesicles were composed of 91% DOPC, 5% DGS-NTA-Ni^2+^, 2% DP-EG10-Biotin, and 2% ATTO 465-DOPE (mol%). Data in **e** and **f** is presented as a 5 nm-increment moving arverage of the raw data, which is composed of >1000 data points. Data in **h** and **i** was binned in increments of 10 nm, rather than 5 nm. Error bars in **e**-**i** represents the standard error of the mean within each bin.

To measure the curvature sensitivity of his-clathrin, we utilized the tethered vesicle assay depicted in Figure 1**b**. Here, polyethylene glycol (PEG) was used to passivate glass cover slips and biotin-NeutrAvidin interactions were used to tether vesicles with diameters ranging from 25-200 nm. This range encompasses the diameter of clathrin coated vesicles, which can vary from 30-100 nm^21, 25^. To promote binding of his-clathrin, DGS-NTA-Ni^2+^ (1,2-dioleyl-sn-glycero-3-[(N-(5-amino-1-carboxypentyl)iminodiacetic acid)succinyl] nickel salt) was incorporated into vesicles at 5 mol%. After the vesicles were tethered, they were incubated with protein-containing solutions. Protein binding was monitored using total internal reflection fluorescence (TIRF) microscopy.

Vesicles were fluorescently labeled with 2 mol% ATTO 465 DOPE (ATTO 465 1,2-dioleoylsn-glycero-3-phosphoethanolamine). Proteins were fluorescently labeled with Alexa Fluor 647, which was conjugated covalently to primary amines within each protein. Using the tethered vesicle assay, we monitored partitioning of proteins between vesicles of different diameters. In addition to his-clathrin, we examined the binding of a negative control protein consisting of the epsin N-terminal homology (ENTH) domain, in which the membrane-binding helix, h0, was replaced with a hexa-histidine tag, his-Δh0ENTH, Figure 1**c**. Because h0 is responsible for curvature sensing by ENTH, his-Δh0ENTH lacks significant sensitivity to membrane curvature, as we have previously demonstrated^13^. In Figure 1**d**, the boxed puncta indicate that the ratio of protein fluorescence intensity to lipid fluorescence intensity did not vary substantially with increasing lipid fluorescence, a proxy for vesicle diameter. Therefore, the vesicles appeared a uniform yellow in color when the red (Alexa Fluor 647-protein) and green (ATTO 465 DOPE-lipid) channels were merged. These comparisons suggest qualitatively that his-Δh0ENTH exhibited negligible sensitivity to membrane curvature, consistent with our previous finding^13^. In contrast, his-clathrin appeared to have significant sensitivity to membrane curvature, as indicated by boxed puncta in Figure 1**d**. Here the ratio of protein fluorescence intensity to lipid fluorescence intensity generally increased as lipid intensity decreased. Therefore, when the protein and lipid fluorescence channels were merged, smaller, less intense, vesicles appeared slightly red, while larger, more intense, vesicles appeared slightly green, Figure 1**d**.

To quantify the curvature sensitivity of his-clathrin, we measured fluorescence intensities of colocalized puncta within the lipid and protein fluorescence channels. Using these intensities, we were able to estimate the diameter of each vesicle and the number of proteins bound to it. The vesicle diameters were estimated by comparing the mean value of the diameter distribution obtained from dynamic light scattering to the mean value of the fluorescence intensity distribution for tethered vesicles, as outlined in the methods section^26^. Using TIRF microscopy, all vesicles with diameters less than 200 nm were illuminated uniformly in the evanescent field, Supplementary Figure S1. The number of proteins bound to each vesicle was estimated by dividing the total protein fluorescence intensity by the calibrated fluorescence intensity of a single Alexa Fluor 647-labeled protein, as validated previously^13, 14^, Supplementary Figure S2. For his-Δh0ENTH and his-clathrin, the number of bound proteins increased monotonically as the vesicle diameter increased, Figure 1**e**. This trend is consistent with our previous results^13, 14^ and is attributed to larger vesicles having greater surface area, thus providing increased capacity for protein binding.

For a curvature sensitive protein, the density of membrane-bound protein is expected to increase as vesicle diameter decreases. Here density is defined as the number of proteins bound per vesicle surface area. Therefore, to evaluate the curvature sensitivity of his-clathrin, we plotted relative density as a function of vesicle diameter, Figure 1**f**. Relative density is defined as the protein density at any given vesicle diameter normalized by the density in a reference diameter range of 160-200 nm. As expected, the relative density of his-Δh0ENTH remained nearly constant at all concentrations observed, increasing only 2-fold as vesicle diameter decreased from the reference range to 25 nm. Interestingly, his-clathrin exhibited a significant degree of curvature sensitivity, having a relative density on 25 nm vesicles that was 14 times higher than its density on the reference vesicles. As the his-clathrin concentration was increased from 1 nM to 100 nM and the membrane surfaces approached saturation, curvature sensitivity for his-clathrin decreased, as expected^27^.

The fractional coverage of the membrane surface by proteins is equivalent to the probability that each protein binding site on the membrane is occupied. Curvature sensitivity is reduced when this probability increases, as illustrated in Figure 1**f**. Therefore, when the curvature sensitivities between different proteins are compared, the fractional coverage on the membrane surface must be held constant^13^. Fractional coverage was estimated by multiplying the protein density, which is the number of bound proteins per vesicle surface area, by the projected area of a single protein onto the membrane surface, A_protein_. We used a value of A_protein_ = 105 nm^2^/triskelion for clathrin, assuming assembly of a lattice ^21^. When we compared his-clathrin, wildtype-ENTH, epsin1, and amphiphysin1 at approximately equal coverage of 2-3%, his-clathrin displayed comparable curvature sensitivity to these known curvature sensing proteins ^13^, Figure 1**g**.

Next, we examined the impact of membrane curvature on protein binding equilibria. Using the data from Figure 1**e**, we generated binding curves in Figures 1**h** and 1**i** and Supplementary Figure S3. These binding curves were fit using a Langmuir adsorption isotherm^28^, Equation 1. In this equation, B represents the average protein density on the vesicles at a corresponding solution concentration [protein]. B_max_ and K_d_ are regression parameters that correspond to the maximum density of membrane-bound proteins and the dissociation constant, respectively.

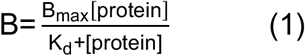

As the solution concentration of his-Δh0ENTH was increased from 5 nM to 300 nM, B/B_max_ reached 80-90%, Figure 1**h** left. The regressed values for K_d_ and B_max_ exhibit no strong correlations with vesicle diameter, as expected, Figure 1**h** right. However, as the solution concentration for his-clathrin was increased from 1 nM to 100 nM, protein density approached saturation much more rapidly, with B/B_max_ reaching 80% at a protein concentration of 20 nM, Figure 1**i** left. As the vesicle diameter increased from 30 nm to 175 nm, the best-fit curves shifted to the right monotonically. This shift corresponds to a monotonic increase in K_d_ with increasing vesicle diameter, Figure 1**i** right. This trend indicates curvature sensitivity, as ln(K_d_) is directly proportional to binding energy^28^. Therefore, the larger the vesicle, the more energy is required for his-clathrin to bind. Notably, the K_d_ values for his-clathrin increased from 1 nM to 5 nM, while the K_d_ values for his-Δh0ENTH fluctuated around an average value of 80 nM. This large difference indicates that his-clathrin had a much stronger affinity for the membrane than his-Δh0ENTH, even though both proteins bind to the membrane using hexa-histidine tags. This increased affinity could be due to the presence of three histidine tags per triskelion or his-clathrin’s ability to assemble into a lattice, which would make binding to the membrane a cooperative process.

To further investigate lattice assembly, we examined the B_max_ values for the most highly curved vesicles in our experiments. Cryo-electron tomography experiments have shown that clathrin coated vesicles contain on average 8-11 triskelia per 1000 nm^2^, about 36 triskelia for a coated vesicle of 30 nm diameter ^21^. In Figure 1i, B_max_ for vesicles of 30 nm diameter reached 9 triskelia per 1000 nm^2^, 25 triskelia in total. This high density suggests that his-clathrin is assembling into lattices on the membrane surface. Specifically, if triskelia remained unassembled, they would occupy a much larger area per molecule, approximately 400 nm^2^ as measured from the crystal structure^16^. This larger area would result in a maximum density of 2-3 triskelia per 1000 nm^2^, around 7 per 30 nm diameter vesicle. A detailed analysis of the dependency of B_max_ on curvature is provided in the supplementary information.

Based on the results from Figure 1, clathrin appears to be a potent sensor of membrane curvature. But what is the mechanism behind this sensitivity? The high curvature sensitivity of scaffolding proteins, such as BAR domains, has previously been attributed to their ability to assemble on membrane surfaces^8, 29^. Therefore, we next probed the curvature sensitivity of his-clathrin under conditions that either enhanced or inhibited its assembly.

### Impact of pH on curvature sensing by clathrin

Clathrin’s ability to assemble depends strongly upon pH^23, 30^. Acidic conditions favor assembly while basic conditions inhibit it. The experiments performed in Figure 1 were carried out at physiological pH 7.4. To perturb assembly conditions, we measured curvature sensitivity at two additional pH values – 6.2 and 8.3. First, we monitored the effect of pH on his-Δh0ENTH binding to see how pH affected the histidine-Ni^2+^ interaction, Figure 2**a**. Three separate sets of wells containing tethered vesicles were each incubated with solutions containing 100 nM his-Δh0ENTH. When compared to the pH 7.4 condition, binding did not change significantly at pH 6.2. However, at an elevated pH of 8.3, binding increased by 40%. This increased binding can be attributed to the deprotonation of histidine, which makes binding to the Ni^2+^ complex more favorable^31^. We then repeated the experiment for his-clathrin with a solution concentration of 2 nM. Here his-clathrin covered approximately 13% of the membrane surface on vesicles with 160-200 nm diameter. Interestingly, his-clathrin exhibited the opposite behavior to his-Δh0ENTH. For his-clathrin, binding increased by 260% at pH 6.2 and decreased by 65% at pH 8.4, Figure 2**a**. This result is consistent with the cooperative nature of his-clathrin’s binding to membranes. At lower pH, where assembly is favored, binding becomes more cooperative as the magnitude of the attractive clathrin-clathrin interactions increases. Although pH 8.3 favored increased binding via the histidine tag, overall binding still decreased. This result suggests that the increased strength of the clathrin-clathrin interactions superseded the decreased strength of the clathrin-membrane interactions.

**Figure 2:**
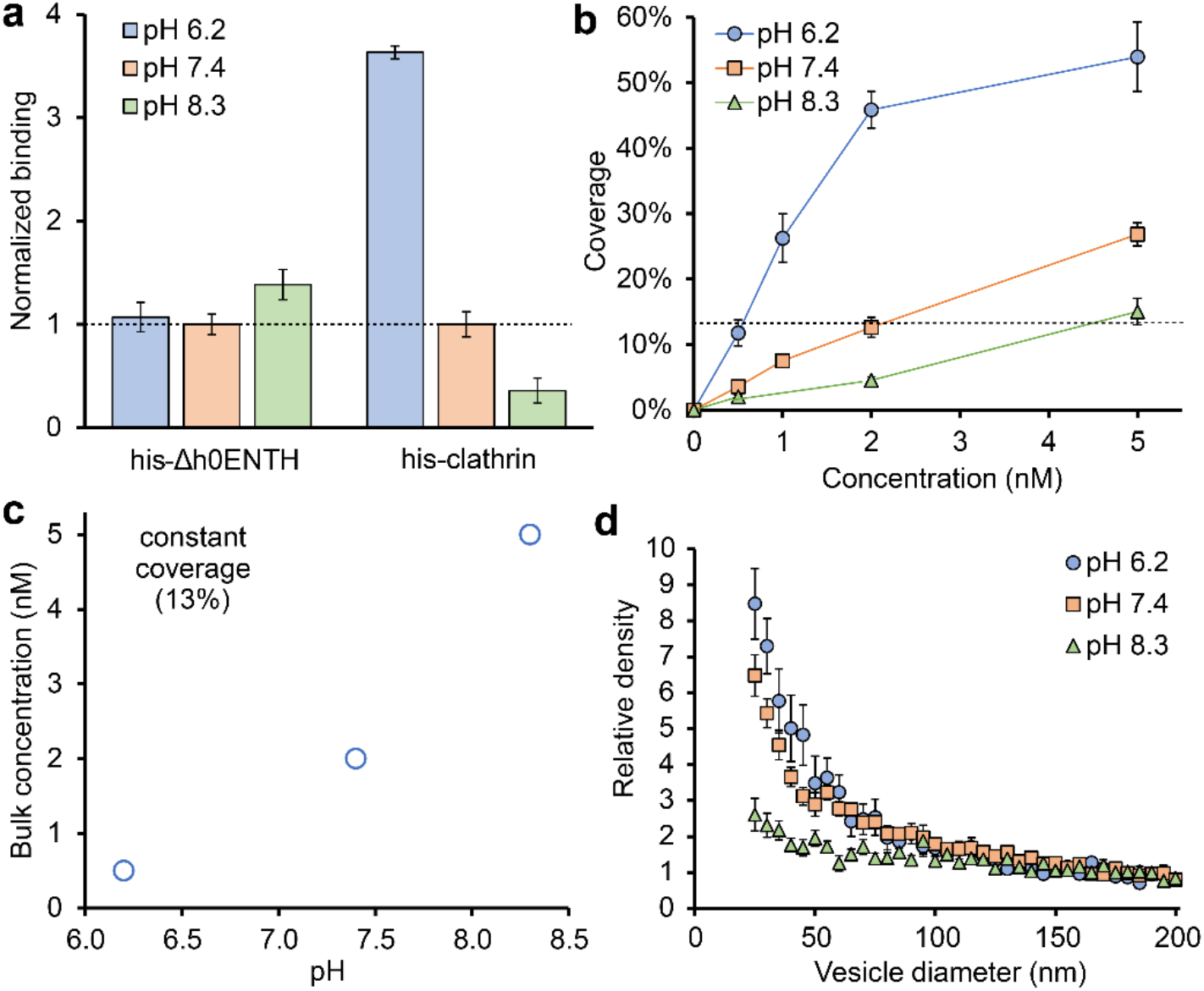
Clathrin’s curvature sensitivity is coupled to its ability to assemble. **(a)** Relative binding for his-Δh0ENTH and his-clathirn on vesicles at different pH values. These quantities were normalized by the coverage values at pH 7.4, which were 2% and 13% for his-ΔENTH and his-clathrin, respectively. The corresponding protein concentrations in solution were 100 nM and 2 nM for his-Δh0ENTH and his-clathrin, respectively. **(b)** Membrane coverage by his-clathrin as a function of solution concentration at various pH values. Solid lines serve connect the data. The dashed line indicates a coverage value of 13%. **(c)** The bulk solution concentrations that yielded 13% coverage on vesicles as a function of pH. **(d)** Relative density of his-clathrin as a function of vesicle diameter. Densities were normalized by the average value for vesicles between 160 and 200 nm. All vesicles were composed of 91% DOPC, 5% DGS-NTA-Ni^2+^, 2% DP-EG10-Biotin, and 2% ATTO 465-DOPE (mol%). Data in **a** and **b** represent average coverage values on vesicles in the 160 to 200 nm diameter range, which was composed of >100 data points. Data in **d** is presented as a 5 nm-increment moving average of the raw data, which is composed of >1000 data points. Error bars in **a**, **b**, and **d** represent the standard error of the mean within each bin.

In Figure 2**b**, we increased the clathrin concentration from 0.5 to 5 nM for the three different pH values. As pH decreased, the amount of clathrin that bound to vesicles increased, consistent with the results in Figure 2**a**. Interestingly, for pH 7.4 and pH 8.3, coverage increased somewhat linearly in the 0.5 – 5 nM range, while coverage at pH 6.2 began to exhibit signs of saturation. These results further suggest that assembly of his-clathrin on the membrane is responsive to changes in pH. In Figure 2**b**, coverages were matched for the three pH conditions at a value of approximately 13% in the reference range. This coverage corresponded to solution concentrations of 0.5 nM, 2 nM, and 5 nM for pH 6.2, pH 7, and pH 8.3, respectively, Figure 2**c**. When we compared curvature sensitivities at these matched coverages in Figure 2**d**, his-clathrin appeared to be most sensitive at pH 6.2, exhibiting a 9-fold increase in partitioning to vesicles with 25 nm diameters. At pH 7.4, his-clathrin density increased 7-fold on small vesicles. Surprisingly, curvature sensitivity at pH 8.3 decreased substantially, exhibiting only a 2-to 3-fold increase in density on vesicles with 25 nm diameters. This low level of curvature sensitivity likely arose from the fractional increase in area for protein binding associated with high curvature, a geometrical effect captured by Supplementary Equation S1. The trend of increasing curvature sensitivity with increasing pH suggests that his-clathrin’s ability to sense membrane curvature arises from its ability to assemble. From this result, we can infer that clathrin triskelia possess preferred angles of interaction, which may become less flexible as lattice assembly precedes^32^, as suggested previously^16^.

### Curvature sensitivity of clathrin recruited by amphiphysin1

We next asked how recruitment by adaptor proteins, which is what occurs during endocytosis, impacts membrane curvature sensing by clathrin. We first investigated clathrin recruitment by the adaptor protein, amphiphysin1. Amphiphysin1 was chosen because it is among the strongest curvature sensors measured to date^13^, Figure 1**g**. It is composed of a 242 residue N-BAR domain, a 380 residue intrinsically disordered domain, and a 73 residue SH3 domain, Figure 3**a**. Amphiphysin1 senses curvature via synergy between the N-BAR domain and the intrinsically disordered region^13, 33^. Within the intrinsically disordered region, there are two motifs that bind to the terminal domain of clathrin: an LLDLD motif at residues 351-355 and a PWDLW motif at residues 380-384^34, 35^.

**Figure 3:**
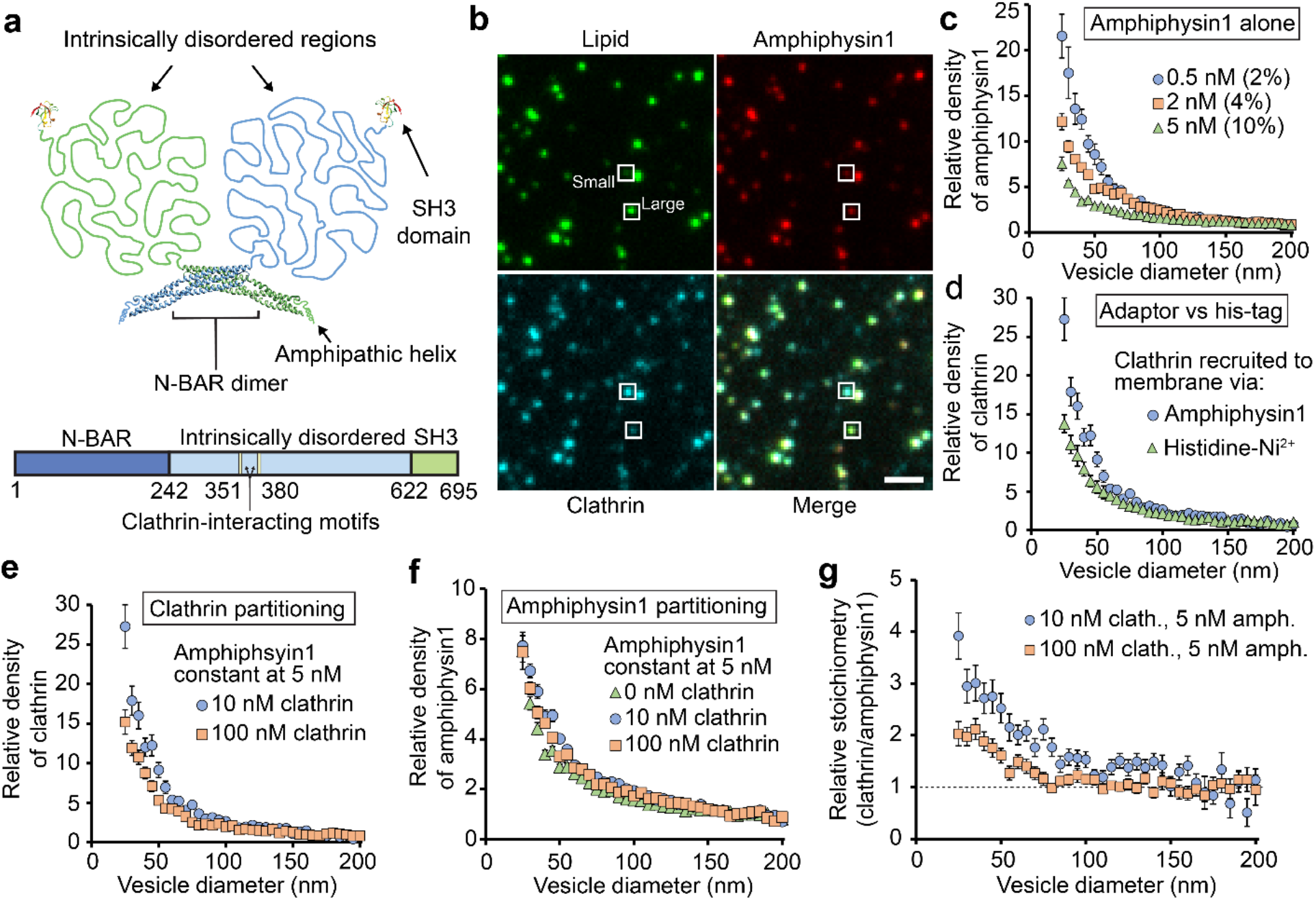
Amphiphysin1 amplifies the curvature sensivity of clathrin. **(a)** Schematic of amphiphysin1’s structure, which includes an N-BAR domain (Protein Data Bank 4ATM), SH3 domain (Protein Data Bank 1BB9), and intrinsically disordered region. **(b)** Representative fluorescent images of tethered vesicles that were incubated simultaneously with amphiphysin1 and clathrin. Squares highlight representative large vesicles. Vesicles were fluorescently labeled with ATTO 465-DOPE. Amphiphysin1 was fluorescenctly labeled with ATTO 594. Clathrin was fluorescently labeled with Alexa fluor 647. Protein concentrations used in these images were 5 nM for amphiphsyin1 and 10 nM for clathrin. Scale bar represents a distance of 2 μm. **(c)** Relative density for amphiphysin1 alone at different solution concentrations. Values in parentheses denote the average fractional coverages in the 160 to 200 nm diameter range. **(d)** Comparison of clathrin curvature sensitivity when recruited to membrane by amphiphysin1 or histidine-Ni2+ interaction (from Figure 1**f**). **(e)** Curvature sensitivity of clathrin in the presence of amphiphysin1. **(f)** Curvature sensitivity of amphiphysin1 at different clathrin concentrations. **(g)** Relative stoichiometry (moles clathrin per mole amphiphysin1) of bound proteins. The absolute stoichiometry was normalized by the average stoichiometry in the 160 nm – 200 nm diameter range. All vesicles were composed of 76% DOPC, 15% DOPS, 5% PI-(4,5)-P2, 2% DP-EG10-Biotin, and 2% ATTO 465-DHPE (mol%). Data in **c**-**f** is presented as a 5 nm-increment moving average of the raw data, which was composed of >1000 data points. Error bars in **c**-**f** represent the standard error of the mean within each bin. Error bars in G represented the propogated error from Supplementary Figure S8.

Using the tethered vesicle assay depicted in Figure 1c, we incubated vesicles (ATTO 465 DOPE) with amphiphysin1 (ATTO 594) and clathrin (Alexa Fluor 647) simultaneously. Fluorescence bleed-through from the ATTO 594 channel to the Alexa Fluor 647 channel was accounted for, Supplementary Figure S4. Figure 3b shows representative fluorescent images of tethered vesicles and the proteins bound to them. There is clear colocalization between the three fluorescent channels. Our experiments utilized an amphiphysin1 oncentration of 5 nM, the minimum for which appreciable levels of clathrin were recruited. Notably, at 5 nM, the curvature sensitivity of amphiphysin1 alone was somewhat reduced from the sensitivity observed at 0.5 nM, owing to increased membrane coverage, Figure 3c.

When the clathrin concentration was 10 nM and the amphiphysin1 concentration was 5 nM, clathrin exhibited a 27-fold increase in relative density on vesicles with 25 nm diameter, compared to a 14-fold increase for clathrin recruited directly to the membrane by its histidine tag, Figure 3**d**. This increased level of curvature sensitivity suggests that adaptor proteins and clathrin work together to enhance curvature sensing. When the clathrin concentration was increased to 100 nM, curvature sensitivity decreased, likely owing to increased coverage^27^, Figure 3**e**. Interestingly, partitioning of amphiphysin1 among vesicles of different sizes is largely unchanged in the presence of clathrin, likely owing to amphiphysin1’s strong affinity for the membrane, which may prevent its repartitioning to more highly curved vesicles^33^. Nonetheless, the ratio of clathrin to amphiphysin1 was higher on small vesicles exposed to either 10 nM or 100 nM clathrin, Figure 3**g**. This finding suggests that clathrin is able to contribute its own sensitivity to curvature on top of the curvature sensitivity of amphiphysin1, helping to explain why clathrin’s curvature sensitivity in the presence of amphiphysin1 exceeds that of amphiphysin1 alone, Figures 3**c** and 3**d**.

### Curvature sensitivity of clathrin recruited by epsin1

Having demonstrated that amphiphysin1 amplifies clathrin’s sensitivity to membrane curvature, we next asked whether epsin1 was able to do the same. Epsin1 is an early-stage clathrin adaptor^36^ and therefore more likely than amphiphysin1 to be involved in recruiting clathrin during endocytosis^37, 38^. Curvature sensing by Epsin1 arises from synergy between the amphipathic helix, h0, within the ENTH domain (residues 1-16) and the intrinsically disordered region (residues 165-575)^13^, Figure 4**a**. Within the intrinsically disordered region, there are two motifs that bind to the terminal domains of clathrin: a LMDLADV motif at residues 257-263 and a LVDLD motif at residues 480-484^35, 39^.

**Figure 4:**
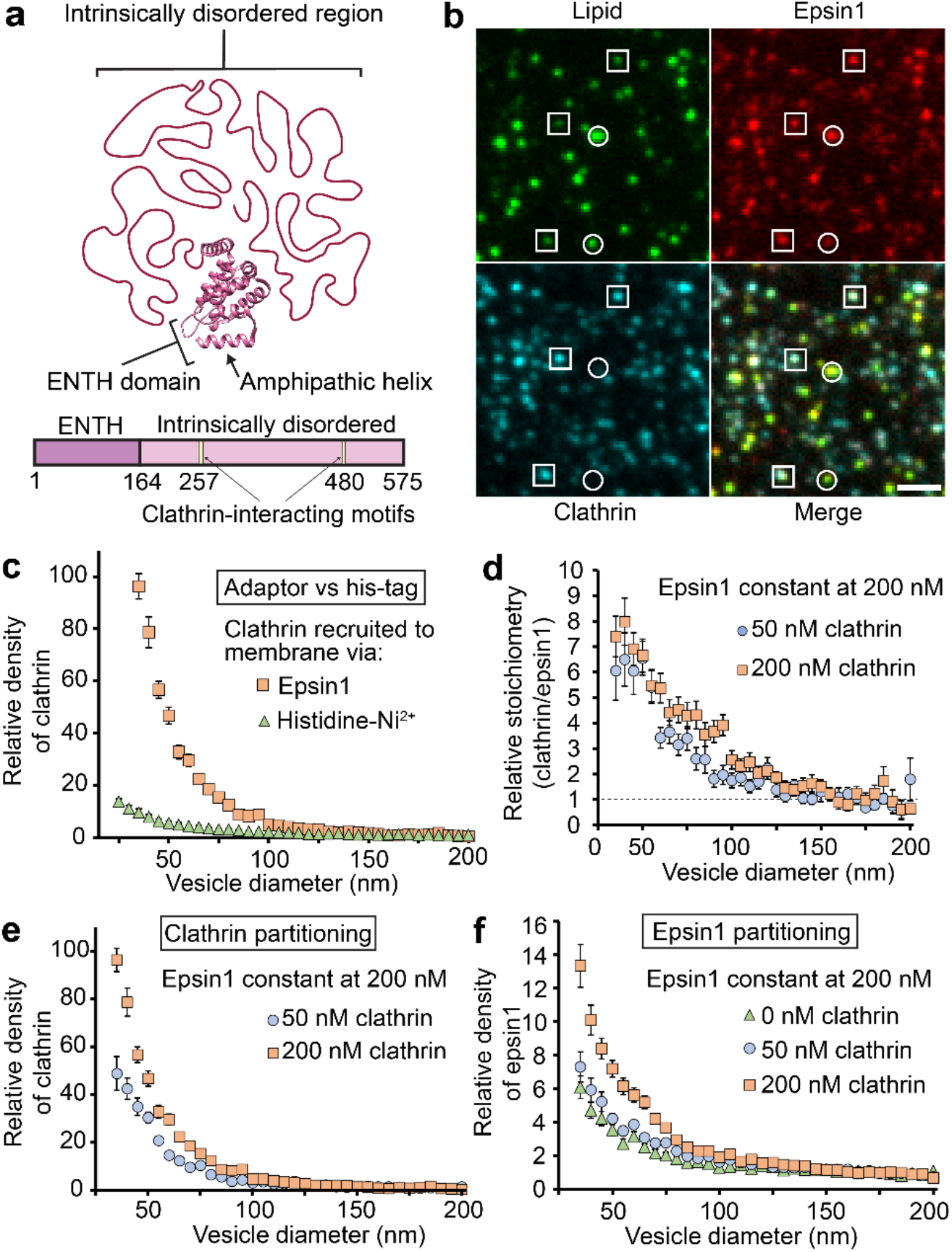
Epsin1 and clathrin cooperatively enhance each other’s sensitivities to membrane curvature. **(a)** Schematic of epsin1’s structure, which includes the ENTH domain (Protein Data Bank 1H0A) and its intrinsically disordered region. **(b)** Representative fluorescent images of tethered vesicles that were incubated simultaneously with epsin1 and clathrin. Squares highlight puncta that are colocalized in all three channels. Circles indicate vesicles that have visible signal in the lipid channel, but no discernable signal in the clathrin channel. Vesicles were fluorescently labeled with ATTO 465-DOPE. Epsin1 was fluorescenctly labeled with ATTO 594. Clathrin was fluorescently labeled with Alexa fluor 647. Protein concentrations used in these images were 200 nM for epsin1 and 50 nM for clathrin. Scale bar represents a distance of 2 μm. **(c)** Clathrin curvature sensitivity when recruited to the membrane by either epsin1 or its histidine tag. **(d)** Curvature sensitivity of clathrin in the presence of epsin1. **(e)** Curvature sensitivity of epsin1 in the presence of clathrin. **(f)** Relative stoichiometry (moles clathrin per mole epsin1) of bound proteins. The absolute stoichiometry was normalized by the average stoichiometry in the 160 nm – 200 nm diameter range. All vesicles were composed of 81% DOPC, 15% PI-(4,5)-P2, 2% DP-EG10-Biotin, and 2% ATTO 465-DHPE (mol%). Data in **c**-**f** is presented as the 5 nm-increment moving average of the raw data, which is composed of <1000 data points. Error bars in **c-e** represent the standard error of the mean within each bin. Error bars in **f** represent the propogated error from Supplementary Figure S9.

In Figure 4**b**, vesicles (ATTO 465-DHPE) were incubated with 200 nM epsin1(ATTO 594) and 50 nM clathrin (Alexa Fluor 647) simultaneously. When the epsin1 concentration was below 200 nM, clathrin was not substantially recruited to the membrane. Above this concentration, most puncta in the lipid channel were colocalized with puncta in the epsin1 and clathrin channels, as indicated by the squares in Figure 4**b**. However, there were also a significant percentage of vesicles that did not have discernable signal in the clathrin channel, as highlighted by circles in Figure 4**b**. In general, these vesicles appeared brighter, and therefore larger, than vesicles that had visible clathrin signal, suggesting that clathrin, when recruited by epsin1, did not bind strongly to vesicles of low curvature, Supplementary Figure S5. A detailed analysis of this result is provided in the supplementary information.

Indeed, when we quantified the partitioning of clathrin in the presence of epsin1, we observed an astonishing 100-fold increase in clathrin density on small vesicles (35 nm diameter) relative to reference vesicles (160-200 nm diameter), Figure 4**c** squares. This level of curvature sensitivity is much greater than the sensitivity observed when clathrin was recruited directly to the membrane by its histidine tag, Figure 4**c** triangles. Furthermore, the curvature sensitivity observed for clathrin in the presence of epsin1 was substantially greater than that of the amphiphysin1-clathrin system, Figure 3, or any protein we have previously studied^13, 14^. What is responsible for the enhanced sensitivity of the epsin1-clathrin system?

Surprisingly, clathrin exhibited an increase in curvature sensitivity when its concentration was increased from 50 nM to 200 nM, Figure 4**d**. This result is the opposite of what we observed for the clathrin-amphiphysin1 system, Figure 3**e**, where clathrin’s curvature sensitivity decreased, likely owing to increased coverage on the membrane surface. Toward explaining this effect, when we examined the partitioning of epsin1 in the presence of clathrin, we observed that epsin1’s curvature sensitivity also increased when the clathrin concentration increased, Figure 4**e**. Specifically, when the clathrin concentration was increased from 0 nM to 200 nM, the relative density of epsin1 on vesicles with 35 nm diameter relative to vesicles in the reference diameter range (160 nm - 200 nm) increased from 6-fold to 13-fold. This result also differs from the amphiphysin1-clathrin system, Figure 3**f**, where amphiphysin1’s distribution among vesicles of different sizes was largely unaffected by clathrin. From these comparisons, we can infer that clathrin and epsin1 work together cooperatively to enhance curvature sensing. Specifically, epsin1 appears able to repartition to more highly curved vesicles in the presence of sufficient densities of clathrin, likely owing to clathrin’s preferential assembly on the surfaces of more highly curved vesicles. This cooperative relationship is further illustrated in Figure 4**f**, which shows that the ratio of clathrin to epsin1 increases with decreasing vesicle diameter. Unlike the clathrin-amphiphysin1 system in Figure 3**g**, this trend becomes slightly stronger, not weaker, as clathrin concentration increases.

## DISCUSSION

Our results demonstrate that clathrin has an inherent capacity for sensing membrane curvature, which grows substantially when it is recruited to the membrane surface by curvature sensitive adaptor proteins. The level of curvature sensitivity observed for clathrin alone is comparable to that of established curvature sensors, such as AP180, epsin1, and amphiphysin1^13, 14, 33^. Further, inhibiting clathrin’s ability to assemble progressively diminished its curvature sensitivity, suggesting that preferred angles of interaction exist within the clathrin lattice. These preferred interactions would likely manifest themselves as optimal arrangements of pentagonal and hexagonal facets within the lattice, as has been suggested based on existing structural data^21, 40^.

When clathrin was recruited to the membrane by the curvature sensitive adaptor proteins amphiphysin1 and epsin1, its curvature sensitivity was substantially amplified. This amplification was more pronounced with epsin1, an early-stage adaptor^36^, than with amphiphysin1, a late-stage adaptor^37, 38^. Furthermore, the epsin1-clathrin system exhibited a high degree of cooperativity, with mutual increases in curvature sensitivity by both epsin1 and clathrin. Clathrin’s lower curvature sensitivity in the presence of amphiphysin1 may arise from amphiphysin1’s high membrane affinity, which makes it more difficult to repartition to highly curved vesicles after clathrin is added. Interestingly, it is amhiphysin1’s ability to form membrane-bound scaffolds that makes it both a strong curvature sensor and a strong membrane binder^33^. Paradoxically, proteins with weaker binding and sensing properties, such as Epsin1, appear better suited to work cooperatively with clathrin, resulting in superior curvature sensing overall. These results are consistent with the idea that curvature sensing is most critical during the early stages of clathrin-mediated endocytosis, when adaptors that bind the membrane relatively weakly, such as epsin1, AP180, and AP2, are responsible for recruiting clathrin^36, 41^.

Our findings may also provide insight toward understanding the mechanism of clathrin-coated pit maturation. In the constant curvature model, endocytic pits maintain a constant radius of curvature throughout their development^18^, whereas in the constant area model, clathrin assembles into a relatively flat lattice that abruptly transitions into a highly curved morphology^19, 20^. Our results show that clathrin is strongly attracted to sites of high membrane curvature. This finding is more consistent with the constant curvature model, in which clathrin stabilizes and extends curved membrane sites. The assembly of clathrin into flat lattices, which is required for the constant area model, appears thermodynamically less favorable. Nonetheless, flat clathrin lattices have been observed in physiological systems^17, 20^. Therefore, future work is needed to explain the counterbalancing forces that overcome clathrin’s inherent preference for curved surfaces, leading to flat lattices. Local increases in membrane tension or receptor density provide possible explanations that warrant further investigation.

Curvature sensing aids membrane remodeling by allowing diverse proteins to partition together into regions of high curvature^42, 43^. Membrane remodeling is essential for membrane trafficking, which includes clathrin-mediated endocytosis^44^. Our findings have revealed that clathrin is not passively recruited to the membrane, but instead possesses strong curvature sensing properties of its own. More broadly, the remarkable increase in curvature sensitivity when clathrin works cooperatively with adaptor proteins helps to explain how multiple proteins within a network can work together to initiate and maintain the curvature of membrane structures throughout the cell.

## METHODS

The methods can be found in the supplementary information index.

## ACKNOWLEDGEMENTS

This research was supported by the National Institutes of Health through R01GM112065 to Stachowiak and Lafer, R01GM118933 to Lafer, and F32GM128316 to Zeno. Purification of epsin1 and histidine-tagged clathrin was performed in the University of Texas Health Science Center at San Antonio Center for Macromolecular Interactions, which is supported by the Mays Center through the National Cancer Institute P30 Grant CA054174, and Texas State funds provided through the UTHSCSA Office of the Vice President for Research.

## AUTHOR CONTRIBUTIONS

All authors designed and performed experiments. In addition, all authors consulted together on the interpretation of results and the preparation of the manuscript.

## COMPETING INTERESTS

The authors declare no competing interests.

## Notes

### Competing Interest Statement

The authors have declared no competing interest.

