## Supporting Information for "Clathrin Senses Membrane Curvature"

[a]Department of Biomedical Engineering, The University of Texas at Austin, Austin, TX 78712; [b]Mork Family Department of Chemical Engineering and Materials Science, University of Southern California, Los Angeles, CA, 90089; [c]Department of Biochemistry and Structural Biology, The University of Texas Health Science Center at San Antonio, San Antonio, TX 78229; [d]Institute for Cellular and Molecular Biology, The University of Texas at Austin, Austin, TX 78712

### SUPPLEMENTARY DISCUSSION

#### Dependence of $B_{max}$ upon curvature for his-clathrin

For his-Δh0ENTH, $B_{max}$ did not vary with diameter substantially and had an average value of 5 proteins per 1000 $nm^2$, Figure 1**h**. Assuming a projected area of 15 $nm^2$ his-Δh0ENTH[1], a density of 5 proteins per 1000 $nm^2$ corresponds to a fractional coverage of 8% on the vesicle surface. This low coverage suggests that the membrane surface was not crowded and is indicative of biochemical saturation, rather than steric saturation. Steric saturation would be expected to occur at a coverage above 30%, where steric pressure between crowded particles becomes significant[2]. If the binding of his-clathrin was also limited by biochemical saturation, it should have saturated the membrane at 1/3 the protein density since it contains 3x the number of histidine-tags. This saturation would be approximately 1.7 proteins per 1000 $nm^2$. However, for his-clathrin, $B_{max}$ increased from 3 to 9 proteins per 1000 $nm^2$ as vesicle diameter decreased from 175 nm to 30 nm, Figure 1**i**. These higher $B_{max}$ values can be attributed to the cooperative binding of his-clathrin to the tethered vesicles. Because his-clathrin can assemble, it was not limited by the number of $Ni^{2+}$ binding sites available on the membrane. Assuming each triskelion has a projected area of 105 $nm^2$ in an assembled clathrin lattice[3], the corresponding fractional coverages for his-clathrin were 32% to 95%. These exceptionally high coverage values strongly suggest that clathrin binding was limited by the assembly of a lattice which covered a substantial portion of the membrane surface.

We sought to determine whether or not this observed dependence of $B_{max}$ on curvature could arise from a geometric effect where highly curved surfaces provide relatively larger area for protein binding, Supplementary Equation S1[4]. Here $r_{clath}$ represents half the height of a single clathrin triskelion bound to the membrane by its terminal domains and $r_{ves}$ represents the radius of the vesicle. Using the available structure for clathrin (Protein Data Bank 3IYV), we measured a value of $r_{clath}$ = 5.8 nm. When we use Supplementary Equation S1 to calculate the relative increase in area for a vesicle with 30 nm diameter relative to a vesicle with 175 nm diameter, we obtain a factor of 1.7. However, in Figure 1**i**, we see that $B_{max}$ increases by a factor of 4 in the 30-175 nm diameter range. Therefore, we can conclude that this observed dependency of $B_{max}$ upon curvature does not cannot be fully accounted for by the geometric effect and likely also arises from clathrin's preferential assembly on smaller vesicles.

$$\frac{\text{available binding area}}{\text{area of vesicle}} = \frac{(r_{ves}+r_{clath})^2}{r_{ves}^2} \qquad \text{(S1)}$$

**Clathrin preferentially binds to small vesicles and avoids large vesicles in the presence of epsin1.**

In Figure 4**b** of the main text, a significant percentage of vesicles did not have discernable signal in the clathrin channel, as highlighted by the white circles. In general, the vesicles lacking significant clathrin fluorescence appeared brighter, and therefore larger, than vesicles that had visible clathrin signal. To rationalize this observation, we defined a fluorescence intensity threshold for the protein channels and examined the distribution of vesicle diameters above and below this threshold, Supplementary Figure S5. This threshold was defined as 3x the standard deviation of the background noise. When we examined vesicles that were colocalized with puncta in the epsin1 channel, we saw that only 9% of vesicles were below the noise threshold and that these vesicles were mostly below 100 nm diameter, Supplementary Figure S5 left. This result can be attributed to poor signal-to-noise ratio for small vesicles, as smaller vesicles tend to have fewer proteins bound and thus have lower signal. Conversely, when we looked at colocalization in the clathrin channel, we saw that 40% of the vesicles had clathrin signal that was below the noise threshold and that these vesicles were mostly above 100 nm diameter, Supplementary Figure S5 right. This result indicates that clathrin was avoiding larger vesicles and mostly binding to smaller ones

### SUPPLEMENTARY METHODS

#### Materials

DOPC (1,2-dioleoyl-sn-glycero-3-phosphocholine), DOPS (1,2-dioleoyl-sn-glycero-3-phospho-L-serine), PI-(4,5)-P2 (L-α-phosphatidylinositol-4,5-bisphosphate ammonium salt) and DGS-NTA-$Ni^{2+}$ (1,2-dioleoyl-sn-glycero-3-[(N-(5-amino-1-carboxypentyl)iminodiacetic acid)succinyl] nickel salt) were purchased from Avanti Polar Lipids, Inc. DP-EG10-biotin (dipalmitoyl-decaethylene-glycol-biotin) was generously provided by Darryl Sasaki from Sandia National Laboratories, Livermore, CA [5]. ATTO 465-DOPE was purchased from ATTO-TEC GmbH. NeutrAvidin, Alexa Fluor 647 NHS Ester, and Zeba spin desalting columns (7K MWCO, 5 mL) were purchased from Thermo Fisher Scientific. TCEP (Tris(2-carboxyethyl) phosphine hydrochloride), PMSF (phenylmethanesulfonyl fluoride), EDTA-free protease inhibitor tablets, imidazole, PLL (poly-L-lysine), ATTO-594 NHS-ester, and Thrombin CleaveCleave Kit were purchased from Sigma-Aldrich. Sodium chloride, HEPES (4-2(2-hydroxymethyl)-1-piparazineethanesulphonic acid), IPTG (isopropyl-β-D-thiogalactopyranoside), BME (β-mercaptoethanol), and Triton X-100 were purchased from Fisher Scientific. Amine reactive PEG (mPEG-Succinimidyl Valerate MW 5000) and PEG-biotin (Biotin-PEG SVA MW 5000) were purchased from Laysan Bio, Inc. Glutathione Sepharose 4B was purchased from GE Healthcare. Amicon Ultra-15 centrifugal filter units were purchased from MilliporeSigma. All reagents were used without additional purification.

**Protein Expression and Purification**

Clathrin: Clathrin heavy chain (CHC) and clathrin light chain (CLC) were expressed and purified as described below utilizing constructs described previously [6]. Immediately before use they were mixed in a 1:1 molar ratio. Even though CLC is purified from a sample of CLC that is co-expressed with CHC, it was difficult to control the stoichiometry, so this method of purifying them separately, quantifying them using densitometry, and reassembling them immediately before use provided a more consistent sample.

Clathrin Heavy Chain (CHC): BL21 competent cells (NEB) were transformed with CHC expression plasmid pET28A (+) 6His-ratCHC-FL, grown in 2xTY + 1/20 volume of 10xM9 salts, 50 μg/ml kanamycin, pH 7.4 at 30$^oC$ to an $OD_{600}$ of 1.4, cooled to 12$^o$ C, induced with 1mM IPTG, and expressed for 24hr at 12$^oC$. Pellets were frozen and stored at -80$^oC$. For purification of CHC, pellets from 2L of cultured cells were defrosted on ice, suspended in 140 mL of 0.5M Tris-HCl, pH 8.0, 5mM TCEP, 1% Triton X-100, 3 tablets EDTA-free protease inhibitor cocktail (Roche), and sonicated at 4 x 2,200 J on ice (OSonica, LLC). The lysate was then batch absorbed to 40 mL bed volume Ni NTA-agarose resin (Qiagen) for 1 hour at 4$^oC$ with stirring at 60 rpm, washed with 10 column volumes 0.4 M Tris-HCL pH 8.0, 1 mM TCEP, 1 mM PMSF, and eluted with 100 mL of 0.5 M Tris-HCl, 200 mM imidazole, pH 8.0, 5 mM TCEP, 2 tablets EDTA-free protease inhibitor cocktail (Roche). The eluate was concentrated by precipitation with an equal volume of saturated ammonium sulfate (in 0.5 M Tris-HCl, pH 7.0), stirred at 60 rpm for 30 minutes at 4$^oC$, collected by centrifugation at 18,000 x g for 40 minutes at 4$^oC$, dissolved in 0.5 M Tris-HCl pH 8.0, 1 mM EDTA, 5 mM DTT, and run on a 330 mL Superose 6 (GE Healthcare) gel filtration column at 4$^oC$. The protein fractions containing CHC were pooled, concentrated by ammonium sulfate precipitation as described above to a final concentration of 10-15 μM, dialyzed into 0.5 M Tris-HCl, pH 8.0, 1 mM EDTA, 5 mM DTT, and the purified CHC was stored as liquid nitrogen pellets at -80$^oC$.

Clathrin Light Chain (CLC): For the purification of CLC, CLC was expressed together with CHC, and the CHC was removed by heat denaturation as follows: BL21 competent cells (NEB) were transformed with CHC expression plasmid pET28A (+) 6His-ratCHC-FL and pBAT4-no tag CLCA1 (co-expressed with pET28A (+) 6His-ratCHC-FL), and the expression and purification were carried out as described above with the addition of 100 μg/ml ampicillin to select for the CLC plasmid, until the resuspension of the first ammonium sulfate pellet. After ammonium sulfate precipitation the sample was resuspended in, and dialyzed against 0.5 M Tris-HCl, pH 8.0, 1 mM EDTA, 5 mM DTT at 4$^oC$ overnight. The next day, the sample was heated in a water bath to 90$^oC$ for 5 min to denature and precipitate the CHC protein which was removed by centrifugation at 8,600 x g for 5 minutes at room temperature. DTT was then added to the CLC to a final concentration of 5 mM, and the purified CLC was stored as liquid nitrogen pellets at -80$^{o}$C.

<u>Epsin1:</u> BL21 Star (DE3) plysS competent cells (Invitrogen) were transformed with epsin1 expression plasmid pGEX6P1-Epsin1FL, grown in 2xYT ,100 µg/ml ampicillin, 34 µg/ml chloramphenicol at 30$^{o}$C to an $OD_{600}$ of 0.8, cooled to 12$^{o}$C, induced with 1 mM IPTG, and expressed for 24hr at 12$^{o}$C. Pellets were frozen and stored at -80$^{o}$C. For purification of epsin1, pellets from 2 L of cultured cells were defrosted on ice, suspended in 120 mL of 0.5 M Tris-HCl, pH 8.0, 5% Glycerol, 5 mM EDTA, 5 mM TCEP, 1% Triton X-100, 3 tablets EDTA-free protease inhibitor cocktail (Roche), and sonicated at 4 x 2,000 J on ice. The lysate was then batch absorbed to 10 mL bed volume of Glutathione Sepharose 4B resin (GE Healthcare) for 1hr at 4$^{o}$C, with stirring at 60 rpm, washed with 10 column volumes of 0.5 M Tris-HCl, pH 8.0, 5% Glycerol, 5 mM EDTA, 1 mM TCEP, 0.2% Triton X-100, 1 mM PMSF, and then washed with 5 column volumes of the wash buffer without Triton X-100. The GST-epsin1 was eluted with 50 mL of 0.5 M Tris-HCl pH 8.0, 5% Glycerol, 5 mM EDTA, 5 mM TCEP, 1 tablet of EDTA-free protease inhibitor cocktail (Roche), 15 mM reduced glutathione. The eluted GST-epsin1 fractions were pooled and concentrated with a 30K MWCO Amicon Ultra - 15 concentrator (Millipore). The buffer was exchanged with Zeba spin desalting columns (Thermo Fisher Scientific) into 50 mM Tris-HCl, pH 8.0, 400 mM NaCl, 5% Glycerol, 5 mM TCEP, and 5 mM EDTA, and the sample was digested with Pierce HRV 3C Protease (Thermo Scientific) for 16hr at 4$^{o}$C. GST and undigested GST-epsin1 were removed by passing the mixture through a 5 mL bed volume Glutathione Sepharose 4B (GE Healthcare) column. The unabsorbed purified epsin1 was then concentrated with a 10K MWCO Amicon Ultra-15 concentrator (Millipore) to a final concentration of 90-100 µM and stored as liquid nitrogen pellets at -80$^{o}$C.

<u>Amphipysin1:</u> Amphiphysin1 were expressed and purified as described previously [4]. DNA coding for residues 2-695 of human Amphiphysin1 was cloned into a pGex4T2 vector, resulting in a construct for a fusion protein with an N-terminal glutathione-S-transferase (GST). The protein was expressed in E. coli BL21 (DE3) pLysS cells. Induction was carried out with 1 mM IPTG at 30°C for 6 hours. The bacteria was then lysed in buffer containing 500 mM Tris, 5 mM EDTA, 10 mM BME, 1 mM PMSF, 5 %v/v glycerol, 1 %v/v Triton X-100, and 1x Roche protease inhibitor cocktail (Sigma-Aldrich) (pH 8.0). Next, bacteria were sonicated on ice (3 × 2000 joules). Bacterial lysate was then separated using ultracentrifugation at 103,800 × g for 40 min. GST-Amphiphsyin1 was then incubated with glutathione agarose beads (Thermo Fisher Scientific). After extensive washing of the glutathione columns, proteins were cleaved directly from the resin by

incubating with soluble HRV-3C protease overnight at 4°C. Excess HRV-3C, which contained a GST tag, was removed by passage through another glutathione agarose column. Amphiphysin1 was them concentrated using Amicon centrifugal filter units (EMD Millipore) and exchanged into buffer containing 25 mM HEPES, 150 mM NaCl, and 5 mM TCEP (pH 7.4) using Zeba Spin Desalting columns. The purified protein was stored at -80°C.

His-Δh0ENTH: His-Δh0ENTH was expressed and purified as described previously [2, 4]. DNA coding for residues 16-164 of rat Epsin1 was cloned into a pRSET vector, resulting in an N-terminal hexa-histidine tag. Following overnight induction with 1 mM IPTG at 18°C, his-Δh0ENTH was then expressed in E. coli BL21 (DE3) pLysS cells. The bacterial lysate was incubated in Ni-NTA agarose beads (Qiagen) in buffer containing 25 mM HEPES, 150 mM NaCl, and 5 mM BME (pH 7.4). After thoroughly washing the resin, his-Δh0ENTH was eluted by incorporating imidazole into the buffer in a stepwise manner up to a final concentration of 200 mM. The eluted protein was then concentrated and dialyzed against the incubation buffer at 4°C overnight to remove imidazole. The purified protein was then stored at -80°C.

**Protein Labeling**
ATTO-594 NHS-ester and Alexa Fluor 647 NHS-ester were dissolved in dimethyl sulfoxide (DMSO) at concentrations of 10 mM and 5 nM, respectively, and stored at -80°C.

Amphiphysin1, Epsin1, and his-Δh0ENTH: Primary amines within these proteins were labeled in buffer consisting of 25 mM MOPS, 150 mM NaCl, and 20 mM BME (pH = 7.4). Protein concentration varied from 50-100 µM. The dye solution was added to 100 µL of the protein solution such that the DMSO never exceeded 1 v/v% and the stoichiometric ratio of dye:protein was 2:1. This mixture was then allowed to react for 30 minutes at room temperature. The resulting labeling ratios for the proteins varied from 0.8 to 1.5 dyes per protein. Unreacted dye was removed using Centri-Spin size exclusion columns (Princeton Separations). Protein and dye concentrations were measured using UV-Vis spectroscopy, and labelled proteins were stored as 5 µL aliquots at -80°C.

Clathrin: Clathrin was freshly labeled prior to each experiment. Clathrin light chain and clathrin heavy chain were thawed on ice and combined at a 1:1 stoichiometric ratio to yield a solution of 3 µM triskelia. Using a Centri-Spin size exclusion column, the clathrin mixture was exchanged into buffer containing 100 mM sodium bicarbonate and 20 mM BME (pH 8.2). The dye solution was added to the clathrin solution at a stoichiometric ratio of 6 dyes per triskelion. This mixture reacted for 20 minutes at room temperature and was then immediately exchanged into a storage buffer containing 10 mM Tris-HCl and 20 mM

BME (pH 8.0). Clathrin was then centrifuged for 10 minutes at 9,000 x g to remove any aggregates. The final concentration of clathrin was 2 μM and the labeling ratios varied from 0.5 to 1 dye per triskelion. Protein and dye concentrations were measured using UV-Vis spectroscopy.

**Preparation of Vesicles**

Vesicles were prepared as described previously [4]. Briefly, lipid aliquots were stored in chloroform-based solvents at -80°C and brought to room temperature prior to use. Lipids were then combined at the appropriate molar ratios, dried under a nitrogen stream, then dried further under vacuum for at least two hours. The lipid film was then hydrated to a lipid concentration of either 100 or 200 μM in buffer containing 25 mM MOPS, 150 mM NaCl, and 20 mM BME (pH 7.4). For formulations containing PI-(4,5)-P2, 0.5 mM EDTA and 0.5 mM EGTA were included in the buffer to prevent aggregation of PI-(4,5)-P2 by divalent metal contamination. Rehydrated lipid suspensions were thoroughly mixed and held at room temperature for 15 minutes before sonication and extrusion. Part of the lipid suspension was sonicated in an ice bath using a probe tip sonicator (Branson Ultrasonics). The remainder of the lipid suspension was extruded through a 100 nm polycarbonate membrane (Whatman plc). The average diameters of the sonicated and extruded vesicles were 50 nm and 115 nm, respectively, as determined by dynamic light scattering.

**Tethering of Vesicles and Protein Binding**

Imaging wells consisted of 1.6 mm thick silicone gaskets (Grace Bio-Labs) with 5 mm diameter circular holes cut into them. These gaskets were placed directly on top of no. 1.5 glass coverslips. Prior to use, gaskets and coverslips were thoroughly cleaned using Hellmanex (Hellma Analytics) and dried under nitrogen. Once assembled, imaging wells were passivated with a buffer solution containing biotinylated PEG as described previously [4]. After 20 minutes of incubation, passivating agent was rinsed from the wells with a sample buffer containing 25 mM MOPS and 150 mM sodium chloride (pH 7.4). Neutravidin was then added to the wells at a concentration of 0.2 mg/mL and allowed to incubate for 10 minutes prior to additional rinsing with sample buffer that contained 20 mM BME. For experiments with vesicles containing PI-(4,5)-P2, imaging wells were rinsed with sample buffer containing 0.5 mM EDTA and 0.5 mM EGTA. Vesicles were then added to the imaging wells at appropriate concentrations. For vesicles containing PI-(4,5)-P2 were, sonicated and extruded vesicles were mixed together at concentrations of 50 nM and 500 nM lipid, respectively. For all other formulations, sonicated and extruded vesicles were mixed together at concentrations of 1 and 2 μM lipid, respectively. After 10 minutes of incubation, excess vesicles were rinsed from the wells with the appropriate sample buffer. For pH dependent experiments, wells were rinsed with sample buffer that was adjusted to either pH 6.2 or pH 8.3. Finally, imaging wells were rinsed with solutions that contained proteins at the appropriate concentration. For experiments using two proteins, proteins were mixed then immediately added to the imaging wells. After addition of protein, the wells were capped with another glass cover slip and allowed to incubate for at least 30 minutes.

**Fluorescence Microscopy**

Vesicles and proteins were visualized using a TIRF microscope consisting of an OLYMPUS IX73 microscope body, a Photometrics Evolve Delta EMCCD camera, and an Olympus 100x 1.4 N.A. Plan-Apochromat oil immersion objective. The microscope was equipped with lasers at wavelengths 473 nm, 532 nm, and 640 nm exciting samples, and a 635 nm laser for autofocusing. We verified that the vesicles were illuminated uniformly within the evanescent wave field, Supplementary Figure S1 After adsorbing carboxyfluorescein beads (50 – 200 nm diameter) to glass cover slips and imaging them with TIRF, we observed that their fluorescence intensities scaled linearly with their volumes. In addition, we compared the fluorescence intensities of tethered vesicles in both TIRF and widefield and observed that their intensity distributions had nearly identical shapes.

**Image Analysis**

For each fluorescence channel, images were acquired in three consecutive time frames (9 images total). The vast majority of vesicles appeared as diffraction-limited puncta. Each of these puncta were fit to two-dimensional Gaussian functions using cmeAnalysis, a particle detection software that is publicly-available [7]. The lipid fluorescent channel served as the reference channel. For this reference channel, the puncta that were included in our analysis needed to meet two requirements – (i) their amplitudes needed to be significantly higher than the local fluorescence background and (ii) they needed to persist in the same location throughout three consecutive frames. After detecting these puncta, cmeAnalysis used an algorithm to find puncta in the fluorescent protein channel(s) that were spatially colocalized with the centroids of the puncta in the reference channel. The radius of this search region was equal to 3x the standard deviation of the Gaussian fit.

For single-protein samples containing ATTO 465-labeled vesicles and Alexa Fluor 647-labeled protein, no significant fluorescence bleed-through was observed in the Alexa Fluor 647 channel. For two-protein samples containing ATTO 465-labeled vesicles, ATTO 594-labled protein, and Alexa Fluor 647-clathrin, no significant fluorescence bleed-through was observed in the ATTO 594 channel. However, fluorescence bleed-through was observed in the Alexa Fluor 647 channel. To correct for this bleed-through, we determined the intensity of ATTO 594 signal that bled through ($I_{647,bleed}$) and subtracted it from the total intensity measured for each puncta ($I_{647,total}$), Supplementary Figure S4. The resulting values corresponded to the true fluorescence intensities of clathrin ($I_{647,clathrin}$), Supplementary Equation S2.

$$I_{647,total} - I_{647,bleed} = I_{647,clathrin} \qquad (S2)$$

The imaging conditions used for calibrating $I_{647,bleed}$ were identical to the imaging conditions used when clathrin (Alexa Fluor 647) was present.

**Calibration of Vesicle Diameter and Number of Bound Proteins**

Vesicle diameters were calibrated using a technique developed initially by the Stamou research group [8, 9] and used in our research group [4, 10]. Briefly, we acquired the distribution of the vesicle diameters using dynamic light scattering and the distribution of fluorescence intensities for tethered vesicles. We then used the means of these distributions to compute a scaling factor between fluorescence intensity and vesicle surface area. Using this scaling factor, we converted vesicle fluorescence intensities to vesicle diameters.

The number of proteins bound to each vesicle was determined using a fluorescence calibration experiment, Supplementary Figure S2. Here we determined the average fluorescent emission from a single protein that was labeled with either ATTO 594 or Alexa Fluor 647. In the calibration measurements, proteins were adsorbed to glass coverslips and appeared as diffraction-limited puncta. These puncta were then imaged in a time series with 2 second intervals over the course of several minutes. During this long exposure, many puncta underwent photobleaching. This photobleaching often occurred over the course of two consecutive image frames. We referred to this sudden photobleaching as single-step photobleaching. Single-step photobleaching most likely occurred for proteins that contained only 1 fluorophore. Therefore, using cmeAnalysis, we tracked the puncta that underwent single-step photobleaching and determined their mean fluorescence intensity values prior to photobleaching, Supplementary Figure S2. Once the single molecule brightness value was determined, we used the same imaging conditions to measure the total brightness of proteins that were colocalized with tethered vesicles. The number of proteins bound was then estimated by dividing this total brightness value by the single molecule brightness value. Raw data for vesicle diameters and the number of bound proteins is shown in Supplementary Figures S6-S9.

### SUPPLEMENTARY FIGURES

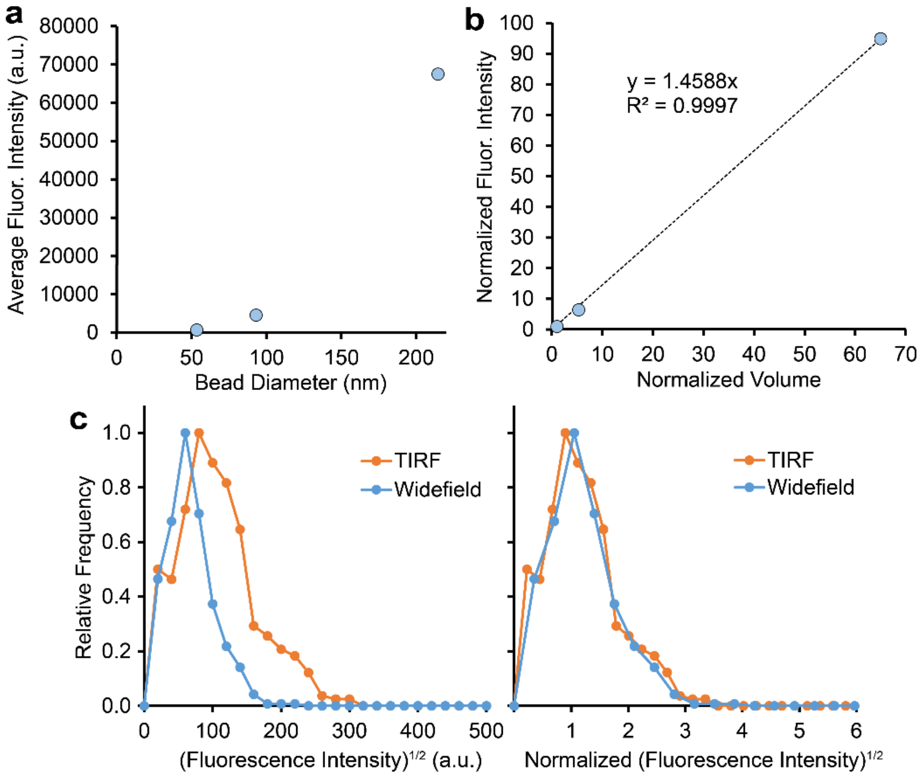


**Supplementary Figure S1. Vesicles were illuminated uniformly with TIRF microscopy. (a)** Average fluorescence intensity of beads that were volume-labeled with carboxyfluorescein. The average diameters of the beads were 53 nm, 93 nm, and 215 nm as measured by dynamic light scattering. These bead populations were incubated separately into three different imaging wells. **(b)** Fluorescence intensity scaled linearly with volume, indicating that the beads were uniformly illuminated in the $\leq$ 215 nm range. **(c)** Comparison of fluorescence intensities for tethered vesicles that were imaged in either TIRF mode or Widefield mode on the microscope. The square roots of intensities were used because (fluorescence intensity)$^{1/2}$ is directly proportional to vesicle diameter. With TIRF imaging, the fluorescence intensities of all objects appeared brighter, as expected. When each distribution was normalized by its mean value, the TIRF and Widefield distributions appeared nearly identical, indicating that vesicles in this size range ($\leq$ 250 nm diameter) were illuminated uniformly.

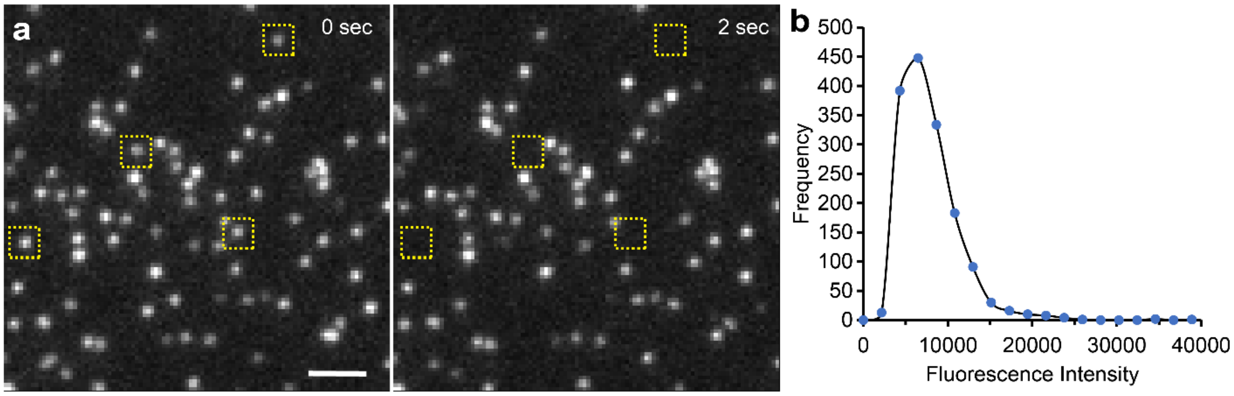


**Supplementary Figure S2. Single molecule calibration of fluorescently labeled protein. (a)** Representative image of fluorescently labeled protein that was adsorbed to a glass coverslip. The boxes highlight puncta that underwent single-step photobleaching. **(b)** Representative fluorescence intensity distribution for puncta prior to single-step photobleaching. This figure depicts his-Δh0ENTH that was fluorescently labeled with Alexa Fluor 647.

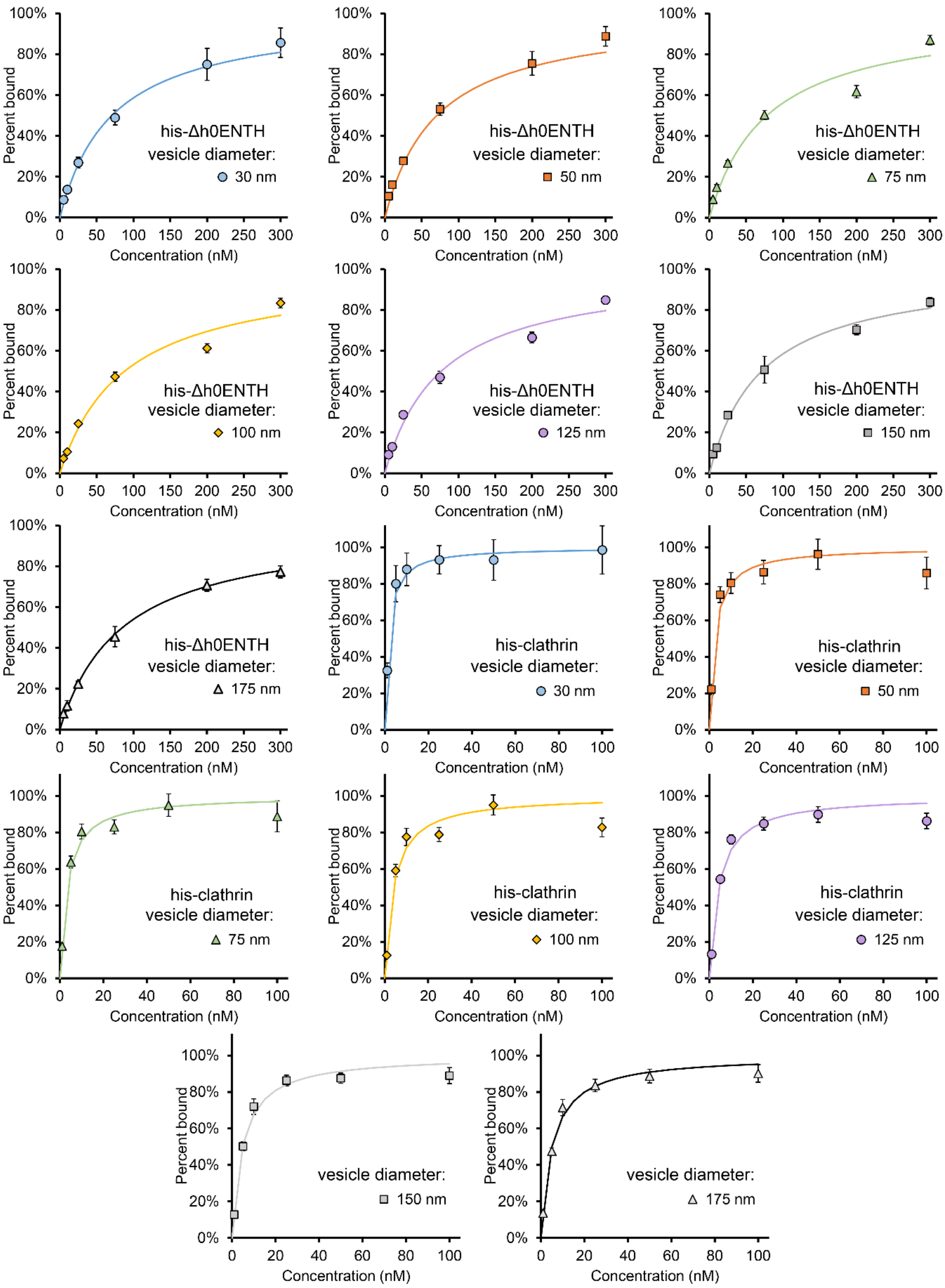


**Supplementary Figure S3.** Individual binding isotherms for his-Δh0ENTH and his-clathrin and the lines of best fit from Equation 1.

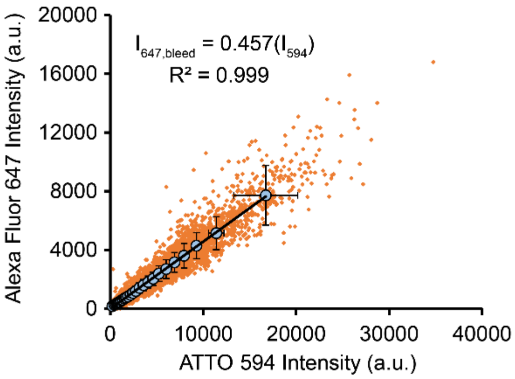


**Supplementary Figure S4. Bleed-through correction for Alexa Fluor 647.** The measured fluorescence intensities of puncta in the Alexa Fluor 647 channel plotted against corresponding fluorescence intensities in the ATTO 594 channel. This representative plot was derived form a sample in which amphiphysin1 (ATTO 594) was bound to tethered vesicles (ATTO 465). Clathrin (Alexa Fluor 647) was not present in this system. For samples in which clathrin was present, $I_{647,bleed}$ was determined for each vesicle from the fluorescence intensity of the colocalized punctum in the ATTO 594 channel ($I_{594}$). The small orange dots represent signal on individual vesicles (N=8042). Blue circles represent the moving average, which was obtained by dividing the data points into 25 equal bins. Error bars correspond to the standard deviation within each bin.

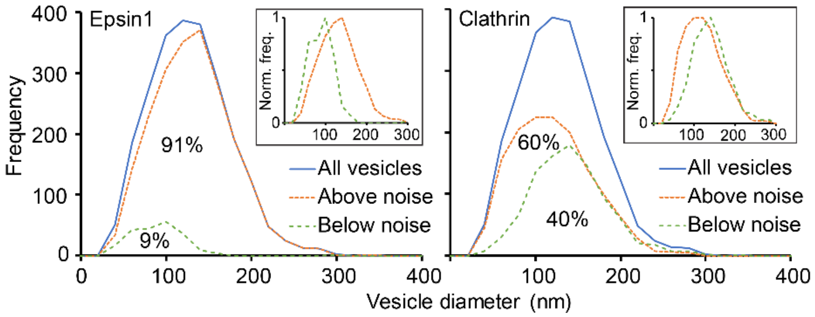


**Supplementary Figure S5. Histograms of vesicle diameters.** Vesicles were split into two populations each for epsin1 and clathrin. These populations consisted of vesicles with signal in the protein channel that was either above or below the background noise. Background noise was defined as 3x the standard deviation of the mean background signal. For these histograms, the concentrations of epsin1 and clathrin were 200 nM and 50 nM, respectively.

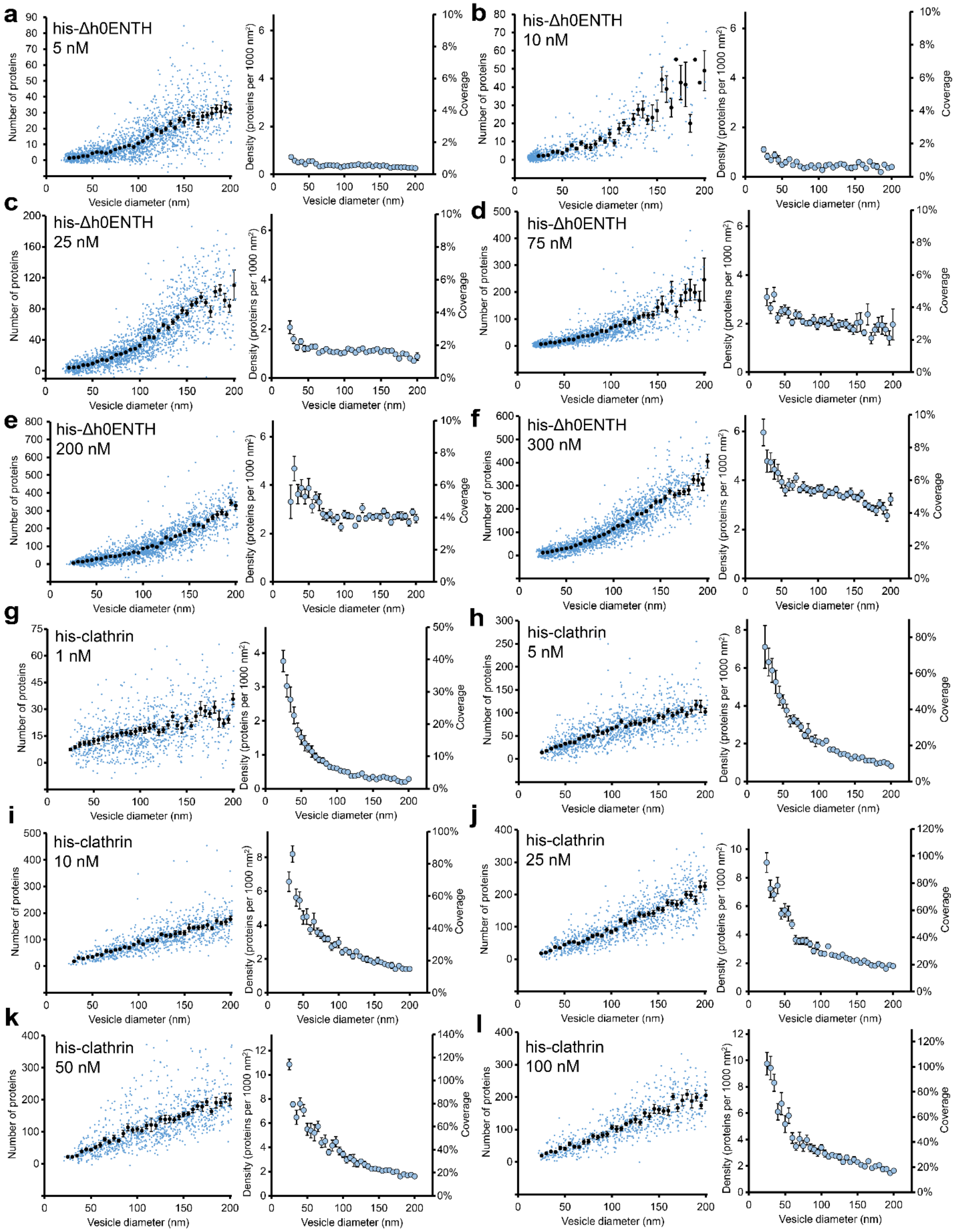


**Supplementary Figure S6. Raw data used to generate Figure 1.** The number of proteins bound with the overlaid moving average in 5-nm increments (left) and corresponding densities and coverages (right) for **(a-f)** his-Δh0ENTH and **(g-l)** his-clathrin. Error bars represent the standard error of the mean within the 5-nm bins.

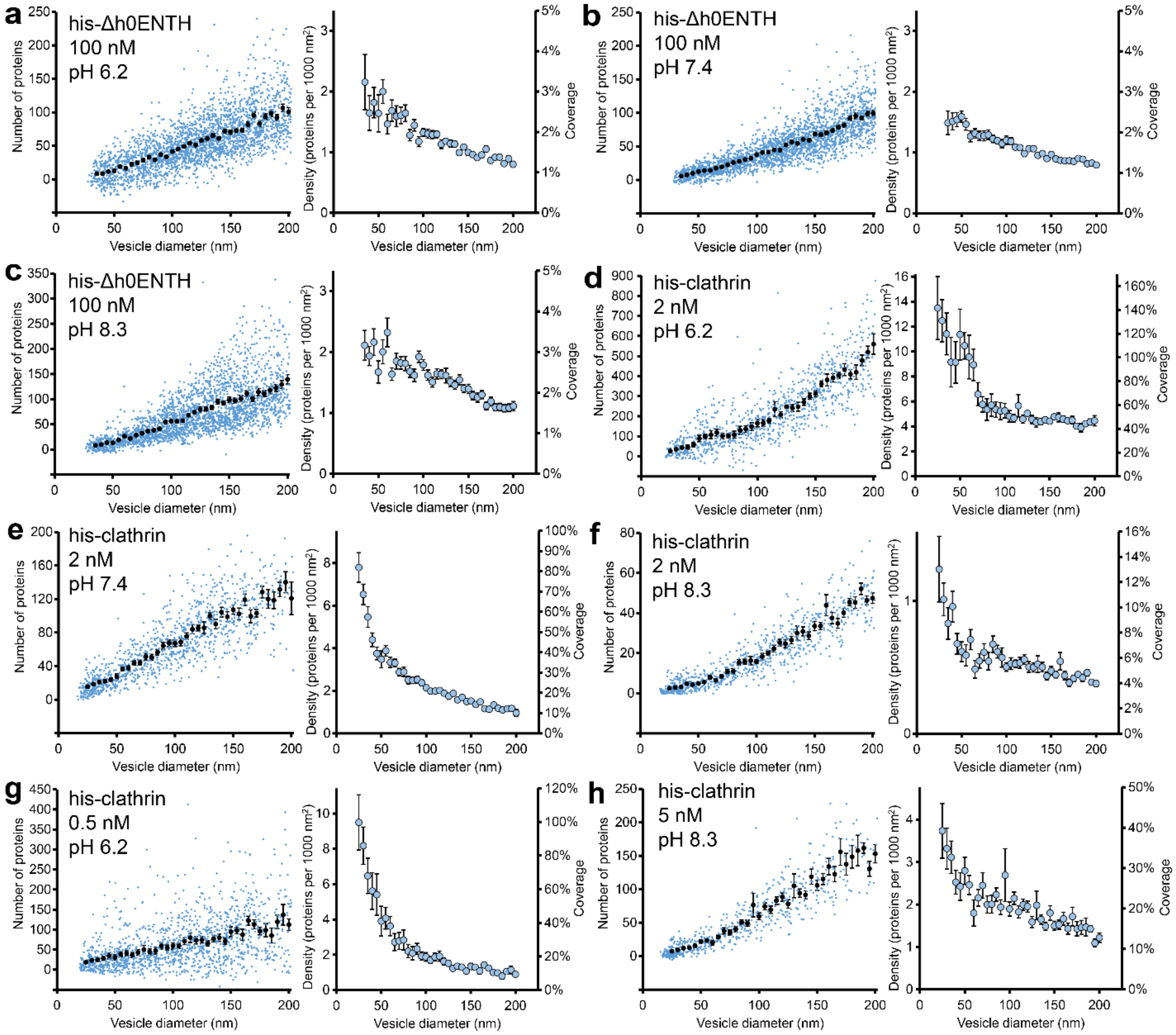


**Supplementary Figure S7. Raw data used to generate Figure 2.** The number of proteins bound with the overlaid moving average in 5-nm increments (left) and corresponding densities and coverages (right) for **(a-c)** his-Δh0ENTH and **(d-h)** his-clathrin. Error bars represent the standard error of the mean within the 5-nm bins.

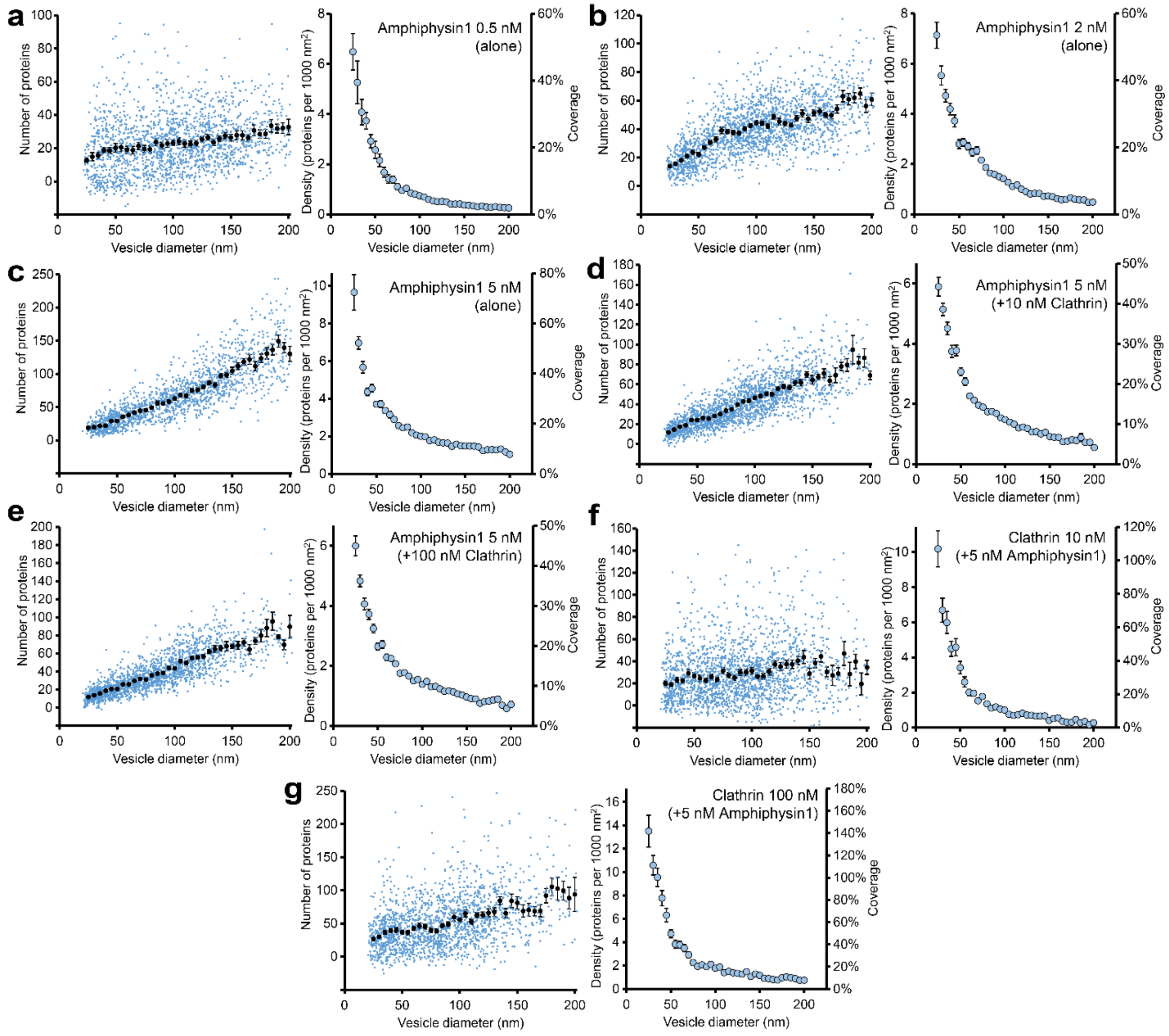


**Supplementary Figure S8. Raw data used to generate Figure 3. (a-g)** The number of proteins bound with the overlaid moving average in 5-nm increments (left) and corresponding densities and coverages (right) for amphiphysin1 and clathrin. Error bars represent the standard error of the mean within the 5-nm bins. A value of $A_{protien}$ = 75 $nm^2$ was used for amphiphysin1.[4]

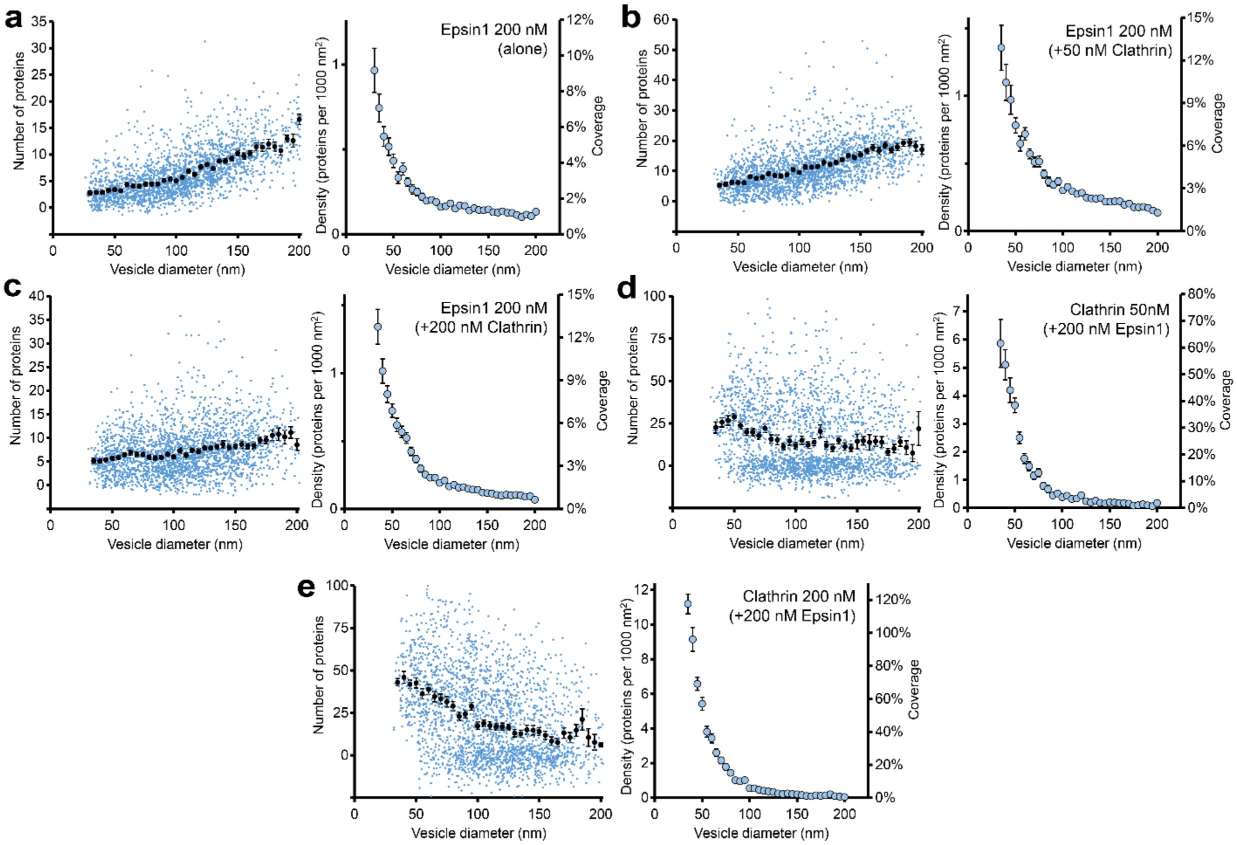


**Supplementary Figure S9. Raw data used to generate Figure 4. (a-e)** The number of proteins bound with the overlaid moving average in 5-nm increments (left) and corresponding densities and coverages (right) for epsin1 and clathrin. Error bars represent the standard error of the mean within the 5-nm bins. A value of $A_{protein} = 95\ nm^2$ was used for epsin1.[4]
